# Excluding Large Grazers Dramatically Improves Survival of Outplanted Juvenile Corals

**DOI:** 10.1101/2024.08.15.607490

**Authors:** Eveline van der Steeg, Adriana Humanes, John C. Bythell, Jamie R. Craggs, Alasdair J. Edwards, Yimnang Golbuu, Liam Lachs, Margaret W. Miller, Janna L. Randle, James R. Guest

## Abstract

High mortality rates of juvenile corals hinder both the natural recovery of populations and the successful implementation of restoration efforts. Grazing is a significant cause of juvenile coral mortality, and grazer exclusion devices have been shown to increase juvenile coral survivorship. However, most experiments have used cages that typically alter water flow and light conditions and exclude grazers of most sizes, making it difficult to quantify the effects of large grazers alone. Here, we test whether deterring large grazers can increase the survival and growth of six-month-old *Acropora digitifera* juveniles outplanted to a shallow reef crest, using arrangements of two or four long or short masonry nails that selectively exclude larger grazers (e.g., parrotfish) while minimising abiotic changes. By the end of our study, colonies with deterrents had significantly larger planar area (almost tenfold for the most effective treatment), more branches, greater height, and enhanced survival than those without deterrents. A critical period is the first week after outplanting when colonies with deterrents had significantly less tissue loss from grazing than those without. Less tissue loss in the first week was associated with significantly higher survival over the following 14 months, with an almost threefold improvement for the most effective treatment. For heavily grazed systems, our study highlights the importance of incorporating grazing deterrents into outplant devices to counteract the negative impact of large grazers on outplanted juvenile coral survival and boost restoration success.

## Introduction

The recovery and resilience of coral reefs depend on successful recruitment after disturbances (Holbrook et al. 2018; Gouezo et al. 2019). High mortality in early life history stages is a critical bottleneck in this process (Babcock 1985; Wilson and Harrison 2005; Miller 2014; Suzuki et al. 2018). The global decline in coral cover (Hughes et al. 2017; Souter et al. 2020) has led to calls for large-scale assisted recovery via restoration and rehabilitation (Anthony et al. 2017). However, restoration success evidenced by outplanted corals reaching reproductive ages has been limited (Boström-Einarsson et al. 2020; Edwards et al. 2024). Asexual fragmentation is currently the most commonly used method to restore degraded reefs (Young et al. 2012; Ferse et al. 2021). However, the low levels of genotypic diversity among fragments, high mortality after outplanting, and challenges in upscaling have been major barriers to restoration success (Omori 2011). The use of sexual reproduction as a method to propagate corals can overcome some of these problems. Substantially higher numbers of outplants with greater genotypic diversity can be produced using such methods, leading to fewer impacts on donor reefs and generating greater upscaling opportunities (Randall et al. 2020; Banaszak et al. 2023). Nonetheless, a major unresolved challenge to implementation is the high level of coral mortality during post-settlement and juvenile stages (Babcock and Mundy 1996; Vermeij and Sandin 2008; Penin et al. 2010; Tebben et al. 2014; Gallagher and Doropoulos 2017). Managing these limitations effectively will determine success or failure within restoration interventions.

Predation and incidental removal by grazing fish can have a considerable negative impact on juvenile coral survivorship (Bak and Engel 1979; Brock 1979; Penin et al. 2010; Lenihan et al. 2011; Doropoulos et al. 2016; Gallagher and Doropoulos 2017). Whether this is due to incidental grazing or targeted feeding to meet dietary requirements (e.g. Clements et al. 2017) is unclear and perhaps varies, but the damage caused and effects on survivorship are well documented (Penin et al. 2010; Doropoulos et al. 2012; Trapon et al. 2013a). Parrotfish are a common culprit, and as larger parrotfish excavate disproportionately more substrate when grazing compared to small fish, they may have a greater impact on juvenile coral survivorship (Bruggemann et al. 1996; Bonaldo and Bellwood 2008; Lokrantz et al. 2008). Despite the negative effects of parrotfish on juvenile corals, these grazers play a crucial ecological role that strengthens coral reef resilience through their suppression of competitive algae (Hawkins and Roberts 2004; Mumby et al. 2006; Shantz et al. 2020). Juvenile coral density and parrotfish biomass are often positively correlated (Hughes et al. 2007; Hoey et al. 2011; Mumby et al. 2013), highlighting the importance of grazers for a healthy ecosystem.

To better understand the factors that determine juvenile coral survivorship, previous studies have focused on physically limiting the access of grazers to corals. However, quantifying the effects of large grazers on juvenile coral survival is challenging as most studies use cages or specially designed substrates that exclude nearly all grazers, which may result in increased algal growth that can compromise coral survivorship. Furthermore, exclusion cages alter abiotic factors, such as water flow, light, and sedimentation, confounding the effect of grazer exclusion. As a result, grazer exclusion studies have found conflicting results, with some showing that grazer exclusion leads to an increase in juvenile coral survival (Baria et al. 2010; Nakamura et al. 2011; Penin et al. 2011; Trapon et al. 2013b), whereas others report a decrease, likely due to intensification of algal growth and competition (Arnold et al. 2010; Steneck et al. 2014; Webster et al. 2015; Doropoulos et al. 2016; Leong et al. 2018). These contradictory results suggest that the effect of grazers on juvenile coral survivorship is highly context-specific and dependent on micro-environmental conditions. For corals outplanted for restoration, adding physical structures that are designed to deter parrotfish instead of excluding them can reduce grazing in the first weeks to months after outplant (Rivas et al. 2021; Pisano 2023; Rule 2023; Whitman et al. 2024), but longer-term effects have not been studied.

Faced with growing stresses and continued declines in coral cover, sexual coral propagation appears to be a promising method for restoring degraded reefs at larger scales. This method can be combined with selective breeding for stress resistance to enhance coral adaptive potential to future climate conditions (Van Oppen et al. 2015; van Oppen et al. 2017). However, outplanted sexually propagated corals face similar survival bottlenecks to natural recruits, with high early mortality being a common occurrence (Omori et al. 2008; Guest et al. 2014; Miller 2014; Chamberland et al. 2017; Humanes et al. 2021). One way to increase outplant survival is to rear corals in nurseries for longer periods to reach an “escape size” prior to outplanting (Guest et al. 2014). However, nursery rearing may not be feasible everywhere as it requires land-based aquarium facilities or suitable locations for ocean nurseries sheltered from storms. If corals can be transplanted to target areas earlier while overcoming juvenile mortality bottlenecks, such as predation, restoration success could increase considerably. Therefore, there is a growing need to design minimally invasive methods to selectively exclude larger grazers that have the potential to harm outplanted corals and compromise their growth and survivorship.

The detrimental effect of large grazers on small juvenile corals has the potential to hinder the efficacy of restoration efforts. Despite this, potential grazing effects on coral survival beyond the first few months after settlement remain poorly understood. To estimate the effect of large grazers on mortality and growth, we outplanted six-month-old sexually propagated juvenile *Acropora digitifera* colonies to the reef with a series of different large grazer deterrent treatments. We estimated the long-term impacts of initial grazing during the first week after outplant and conducted video assays to identify potential grazers. Overall, we found a positive impact of grazing deterrents on juvenile coral survival and growth, indicating that the development of low-cost devices could boost the success of coral restoration efforts.

## Methods

### Study site

This study was conducted in the Republic of Palau in the western Pacific Ocean. Coral spawning and larval rearing were carried out at the Palau International Coral Reef Center (PICRC) in April 2020. The source and outplant site chosen for this experiment is a protected outer reef in the east of Palau (Mascherchur, N 07°17’29.9”; E 134°31’08.0”). The site was selected for its proximity to PICRC and the high abundance of *Acropora digitifera* at 0.5 to 4 m depth. All research was conducted under National Marine Research Permits RE-20-04 and RE-21-02.

### Coral rearing

On April 3^rd^, 2020, five days before the full moon, ten gravid (i.e., containing visible pigmented oocytes) *A. digitifera* colonies with diameters >20 cm were collected on scuba using a hammer and chisel. Colonies were transported to PICRC and maintained in a 760 L shaded outdoor holding tank. The tank was provided with a continuous flow of 50 µm filtered seawater (FSW) and two magnetic pumps, both attached to three flow accelerators to ensure water movement (Pondmaster 1200 GPH, Accel Aquatic Vortex). The tank was lit with four LED aquarium lights (122 cm Reef Bright XHO 50/50). After sunset (19:00 h), the inflow and pumps were turned off, and the tank was covered to block incident light. On April 6^th^, 2020, two days before the full moon, setting was observed in three colonies at 19:45, and they spawned at 20:30. Coral gamete collection and larval rearing followed standard coral propagation methods (Guest et al. 2010, Online Resources).

Larvae were offered previously conditioned settlement units (SUs) in two 45 L settlement tanks, each containing 500 SUs, at a density of 25 larvae per SU. Ceramic SUs (Ocean Wonders LLC, 19 mm in diameter with a 17 mm long stem) were biologically conditioned with fresh crustose coralline algae (CCA) fragments collected from the study site in four 180 L flow-through tanks each with two pumps (Hydor Koralia Nano 240 Circulation Pump/Powerhead) for six months prior to the experiment. Fragments from the parental colonies were added to the tanks to provide a source of Symbiodiniaceae for newly settled corals. Eleven days after larvae were introduced to SUs, the upper surfaces of 26 SUs per settlement tank were examined under a dissecting microscope to count the number of settled corals. The mean settlement rate was 11.3 ± 0.8 corals per SU (4.0 ± 0.3 per cm^2^), resulting in a settlement success of 45 ± 3.0 % (mean ± standard error). Most settled SUs (91%) had at least one live coral after approximately five months (162 days) in the *ex-situ* nursery. Numerous grazers were added to each tank to control algal growth (i.e. juvenile rabbitfish, cerithid snails, trochus snails and small filefish), and nursery tanks were cleaned biweekly. Corals were fed thrice weekly with 0.75 gr Aqua Core Coral Fusion plankton powder.

### Coral outplant with grazing deterrents

Experimental treatments were designed to exclude larger grazers (grazing fish > 20 cm Lokrantz et al. 2008) while allowing access to smaller algae-grazing fish and minimising alteration of abiotic factors such as water flow, light and sedimentation. Six months (185 days; Oct 2020) after settlement, the SUs with the largest juvenile colonies were selected (n = 200 colonies); these had a mean diameter of 2.2 ± 0.25 cm and planar area of 3.88 ± 0.87 cm^2^. These colonies were outplanted to Mascherchur Reef at 20 cm intervals along four 10 m transects, resulting in 50 corals per transect. To attach each colony, a hole was drilled into the bare reef substrate using a battery-powered submersible drill (Nemo Divers Drill) with an 11 mm drill bit, the substrate brushed clean of algae, and the stems of the SUs secured into the drilled holes with epoxy (Milliput Standard). Using nails arranged around the outplanted corals, five grazing deterrent treatments were applied to the outplants: a) 4 long (10.1 cm) masonry nails, b) 4 short (7.6 cm) nails, c) 2 long nails, d) 2 short nails and e) a control with no deterrents (n = 40 per treatment, Fig. 1), hereafter referred to as 4L, 4S, 2L, 2S and control respectively. The mean protrusion of long and short nails after attachment to the substrate was 7.1 ± 0.9 cm and 5.1 ± 0.7 cm, respectively. The mean distance between the heads of the nails (measured diagonally for the four nail treatments) was 5.3 ± 2.1 cm.

**Fig. 1.**
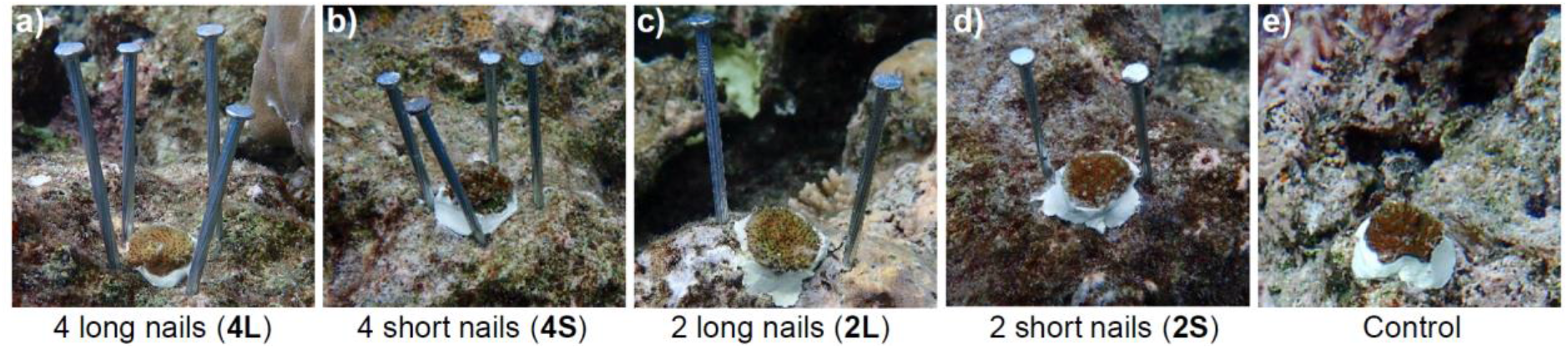
Representative photos of the different levels of large grazer deterrent treatments added to juvenile coral colonies outplanted six months after settlement. The colonies had a mean diameter of 2.2 ± 0.25 cm.

SUs were surveyed at 7, 35, 62, 98, 140, 160, 227, 341 and 425 days after outplant. During surveys, the status of each coral was classified as either alive, missing (SU missing or not found) or dead (entire coral removed from the SU, e.g., Fig. 4e, or intact skeleton without tissue) and any missing nails were replaced. The nails became overgrown with CCA and turf algae through the experiment but remained free from macroalgae. Colony planar area was measured from photos taken during the outplant, 7- and 425-day surveys using ImageJ’s (1.48v, National Institutes of Health, USA). During the last survey, the height of each colony was also measured with callipers and the number of branches was counted.

### Fish grazing estimation

In an attempt to assess the reduction in grazing that resulted from adding grazing deterrents, grazing pressure was estimated with video surveys. Three additional sets of five SUs with corals were outplanted in April 2022. Corals from the same cohort with sizes comparable to the initial size of outplants (∼2.2 cm diameter) were used. They had been maintained in nursery tanks for 18 months, moved to *in-situ* nurseries for five months where they were grazed and reduced in size, and taken back to the *ex-situ* nursery for one month before the outplant. The *in-situ* nursery was located 2.2 km to the west of the study site on a sheltered sand patch surrounded by arborescent *Acropora* thickets (N07°18’19.8’’; E134°30’6.70’’, described in Humanes et al. 2021). SUs were outplanted approximately 20 cm apart, and the treatment order was randomised, with a minimum distance of 20 m between each set. GoPro cameras were deployed one to three times per day between 9:00 and 16:00 on days three, four and five after outplanting. No divers were in the area after deployment, and the first three minutes of each recording were disregarded to eliminate the effect of camera deployment on fish behaviour. This resulted in 21 videos (49 to 116 minutes long) and a total of 27 hours 40 min of footage. Every fish observed in the recordings that took a bite directly on a coral outplant was identified to species, its length was estimated to the nearest 5 cm category (5, 10, 15 or 20 cm), and the total number of bites per treatment was recorded.

### Data analysis

Results are given as mean values ± standard error of the mean. Differences in survival curves between treatments were compared using Kaplan-Meier survival analysis with right-censored data (Lee and Wang 2003). This method is robust for non-normally distributed survival data and allows the inclusion of individuals we could not relocate in the form of right-censored data. When a coral was found to be entirely dead in a survey, the date the coral died was estimated as the midpoint between surveys, as the exact time of death cannot be determined. Log-rank pairwise comparisons were performed to test for significant differences in survival curves. Planar colony area, height and number of branches were compared between treatments with a nonparametric Kruskal-Wallis test followed by a post-hoc Bonferroni corrected pairwise Wilcoxon rank-sum test. The percentage area reduction from grazing in the first week was compared between treatments with a one-way analysis of variance and Tukey’s HSD test. For this analysis, dead corals were included as having an area of 0 cm^2^ (i.e., 100% reduction). The effect of initial grazing intensity (percentage planar colony area reduction in the first week) on survival after 14 months was tested using a generalised linear mixed effect model (GLMM) with a binominal error distribution, with treatments and transects as random effects. To analyse the effect of treatments on bite rate, the bite rates of different fishes were grouped and adjusted to the number of bites per hour and rounded to the nearest whole number. The effect of each grazing deterrent on fish bite rates was tested using a GLMM with a negative binomial error distribution, as data distribution was over-dispersed when first run with a Poisson error distribution. Pairwise comparisons among treatments were conducted using Tukey tests with the “emmeans” R package. All analyses were performed in R 4.2.1 using Rstudio 554 (Rstudio Team 2010; R Core Team 2022).

## Results

### Survival

The mean survival time of corals within all grazer deterrent treatments, except for 2L, was significantly higher compared to the control after 14 months (425 days) on the reef (Fig. 2, Online Resource). The 4L treatment also had a significantly longer survival time than the 2L treatment. The mean survival time of corals outplanted to the reef with the 4L grazing deterrents was 374 ± 18 days, 1.8 times longer than the control 212 ± 28 days. The 4L treatment had the highest survivorship after 14 months, with 65% of the colonies alive; this was almost three times higher than the control, with only 22.5% still alive. On April 18^th^, 2021 (192 days after outplant), typhoon Surigae passed north of Palau, causing high wave energy at the study site. Before typhoon Surigae, at approximately one year old (160 days on the reef), 92% of outplants in the 4L treatment were still alive. Of those colonies still alive one month before typhoon Surigae, 22% had died or were missing one month after the typhoon.

**Fig. 2.**
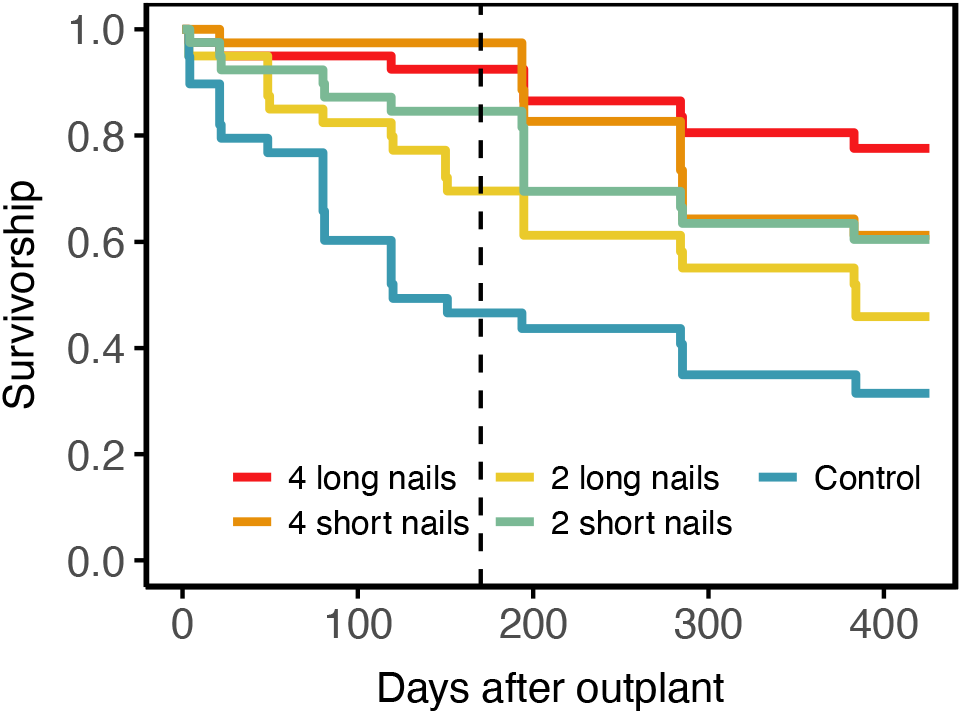
Survival curves of *Acropora digitifera* juvenile colonies after outplanting to the reef at six months old with five different levels of grazing deterrents. The dashed line indicates the timing of Typhoon Surigae.

After 14 months on the reef, the planar area of colonies in both four nail treatments was almost tenfold larger than controls (P < 0.05), while colonies in the two nail treatments had intermediate planar areas (Fig. 3a & b). The mean planar area at the final survey was 13.3 ± 2.5 and 11.7 ± 2.7 cm^2^ for the 4L and 4S treatments, respectively (approximately 3.8 ± 0.8 and 3.4 ± 1.0 cm in diameter). Similarly, the mean number of branches and mean colony height were significantly higher in both four nail treatments compared to the control (Fig. 3c & d). The percentage of live colonies self-attached to the substrate (i.e., coral tissue growing directly on the reef substrate) was 63, 55, 40, 43 and 15% for the 4L, 4S, 2L, 2S and control treatments, respectively.

**Fig. 3.**
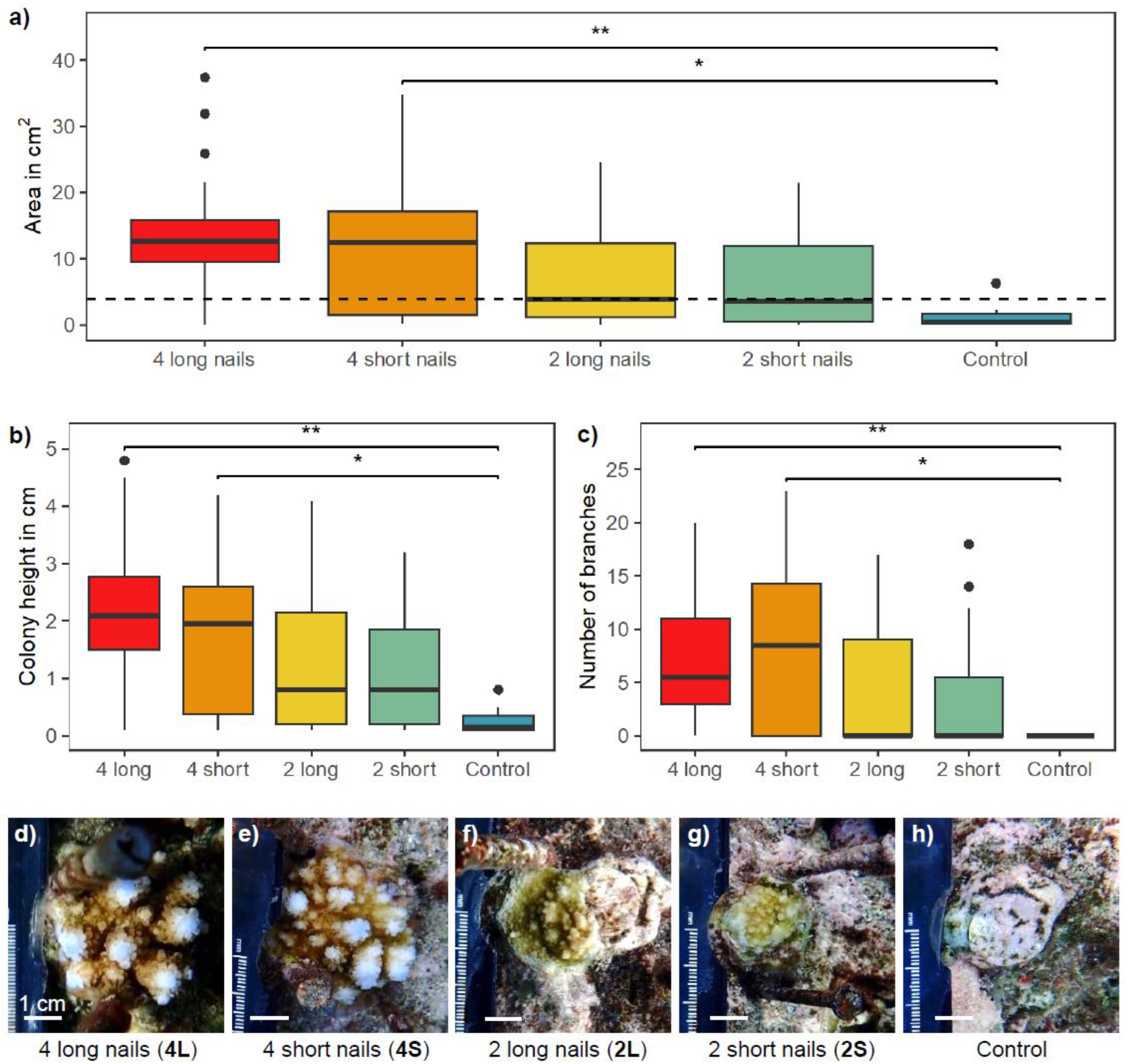
Boxplots of **a)** planar area in cm^2^, **b)** colony height and **c)** number of branches of *Acropora digitifera* juvenile corals outplanted at age six months under five different levels of grazing deterrence after 14 months on the reef. The dashed line in 3a is the mean size of all colonies at time of outplant. Asterisks indicate level of significance between grazing deterrent treatments based on nonparametric Kruskal-Wallis tests and Bonferroni corrected pairwise Wilcoxon rank-sum tests (** *P* <0.01, * < 0.05). **d-h)** Photos of representative colonies with areas close to the median size of each treatment, with scale bars of 1 cm at the end of the study.

**Fig. 4.**
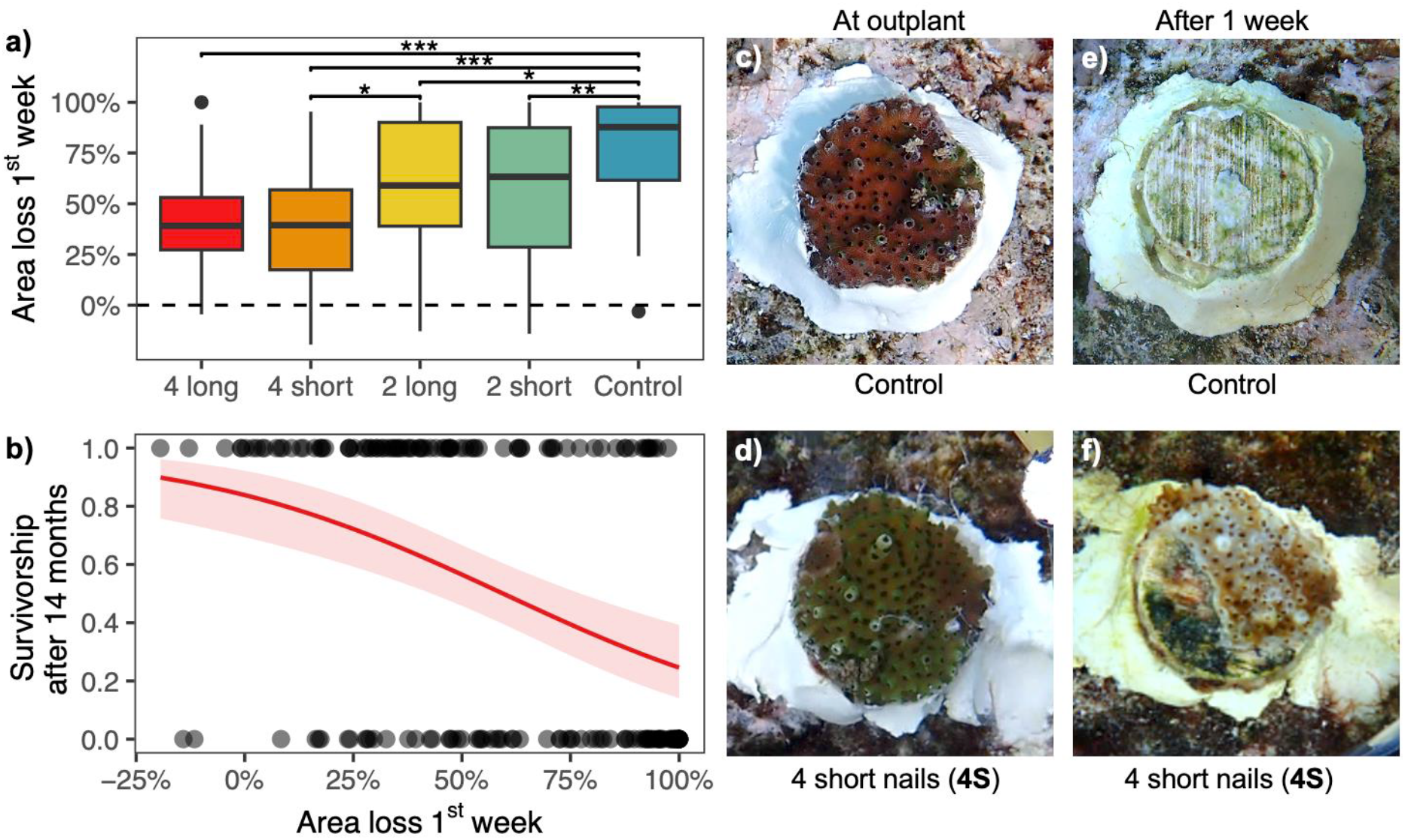
**a)** Percentage loss of planar colony area within one week after outplant, asterisks indicate the level of significance between grazing deterrent treatments based on a one-way ANOVA and Tukey’s HSD test (*** *P* <0.001, ** <0.01, * < 0.05). The dashed line indicates 0% area loss. **b)** The predicted relationship between percentage loss of planar colony area within one week after outplant and survivorship after 14 months on the reef from a binominal GLMM. The shaded area indicates the confidence interval. **c)** Colony from the control treatment with no grazing deterrents at outplant and **e)** the same colony after one week on the reef, grazing scars are visible on the settlement unit, and the coral is missing. **d)** Colony from the 4 short nail treatment at outplant and **f)** the same colony after one week on the reef with part of the colony missing.

### Grazing in the first week post-outplant

There was a large effect of grazing on planar colony area within the first week after the colonies were outplanted (Fig. 4a). Colonies without any grazing deterrents (controls) had a mean reduction in size of 78 ± 4%, which was significantly higher than all other treatments (Fig. 4c & e). There was significantly less reduction in planar area in the 4S treatment compared to the 2L treatment. The 4L treatment also showed an improvement on the 2L treatment but with uncertainty exceeding the alpha threshold of 0.05 (*P* = 0.052). Nonetheless, all treatments suffered a considerable reduction in mean area in the first week, with even corals in the four nail treatments losing up to 40% of tissue area (Fig. 4d & f). Across all treatments, there was a negative effect of initial grazing on long-term survival (Fig. 4b, GLMM, *Z* = -4.3, *P* < 0.001).

### Fish grazing estimation from video

A total of 77 grazing events (a sequence of consecutive bites by a single fish without the fish leaving the frame) were recorded, with 290 individual bites by 14 species, including two parrotfish (*Chlorurus sordidus* and *Scarus psittacus*, Online Resources). There was a significant difference in bite rates between the 4S treatment and both the 2L treatment and control (Fig. 5, Tukey test, *P* < 0.05 for both). No schools of grazing fish were observed around the outplants, and no fish over 20 cm were seen grazing on the colonies. Only one fish of ∼20 cm was observed grazing on a colony in the control treatment and all other fishes were ∼15 cm or smaller. No grazing scars were observed on the colonies after the three days of recording.

**Fig. 5.**
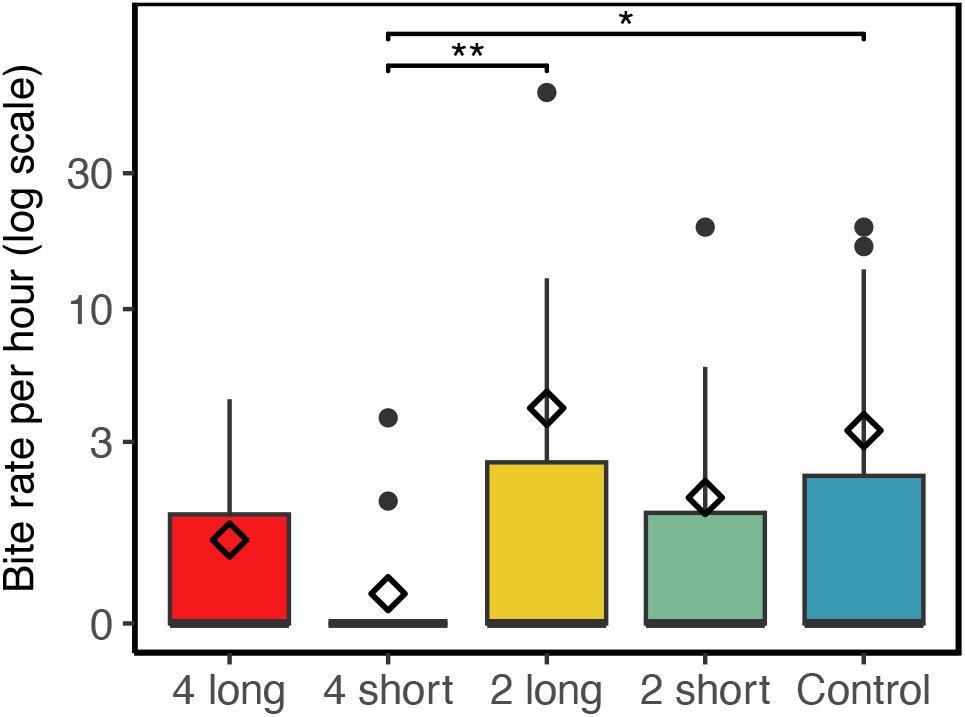
Bite rate recorded as the number of bites per hour of all fishes grouped, tested using a negative binomial generalised linear model paired with a Tukey test for pairwise comparisons, plotted on a log scale. Diamonds represent the mean bite rate per hour. Asterisks indicate level of significance between grazing deterrent treatments (** *P* <0.01, * < 0.05).

## Discussion

Survival bottlenecks in the early ontogeny of corals can hinder both the recovery of wild populations following disturbances and the success of coral restoration efforts. Grazing is known to be a potential moderator of these juvenile survival bottlenecks, as scraping can damage small, settled corals. However, the specific role of large grazers on juvenile coral growth and survival remains poorly understood. Using a manipulative field experiment, we found that excluding large grazers significantly increased juvenile coral survival and growth. Grazing impacts within the first week after outplanting to the reef caused an immediate bottleneck in survivorship and growth for those juvenile corals affected, with lasting effects on the long-term survivorship of even those corals that were still alive after one week. For heavily grazed reefs, our results suggest that simple devices that deter large grazers from grazing newly outplanted corals could mitigate juvenile survival bottlenecks and increase the success of restoration efforts.

In our study, juvenile corals suffered from tissue loss and subsequent mortality, particularly in the first week after outplanting (Fig. 4). Clear evidence of grazing scars on outplanted substrates suggests that the most plausible explanation for this is grazing by large parrotfish. Parrotfish are abundant in Palau, with regular sightings of bumphead parrotfish schools at the outplanting site (Muller-Karanassos et al. 2021; Friedlander et al. 2023; Karanassos et al. 2023). Large parrotfish grazers are known to consume live corals, leaving distinctive grazing scars (Bonaldo and Bellwood 2009; Bonaldo and Rotjan 2018). After observing grazing scars and tissue loss of the corals, video assays were set up on a new set of outplanted colonies to identify the types of fish responsible for the damage. The assays showed a significantly lower mean bite rate in the four short nail grazing deterrent treatment, indicating this treatment was effective at deterring grazers. In addition, small fish were able to access and graze on coral colonies with grazing deterrents without leaving visible grazing scars. However, the number of grazing observations in the assays was limited, and no large parrotfish were captured grazing the corals during these relatively short video survey periods. It is likely that the duration of the video assays was not long enough to capture sporadic intensive grazing events by large groups of parrotfish that could be a major driver of the juvenile survival bottleneck immediately after outplanting.

A diverse range of grazing, scraping, and corallivorous organisms are associated with coral reefs. Although large parrotfish are the most common grazers on tropical reefs, triggerfish have also been observed grazing on coral (Bellwood et al. 2006; Gallagher and Doropoulos 2017), along with crown-of-thorns starfish, which can be devastating to coral populations (Kayal et al. 2012). In addition, nocturnal invertebrates, such as urchins, are known to graze juvenile corals (Bak and van Eys 1975; Glynn 1996; Korzen et al. 2011; O’Leary et al. 2013). This complicates the survey logistics required to identify the taxa affecting colony growth and survivorship. However, physical grazing deterrents are likely effective for multiple grazer and predator groups. Moreover, reefs with different characteristics, such as high sedimentation or algae cover, likely have additional factors affecting juvenile coral survivorship. Thus, site-specific trials should be conducted before large-scale coral outplantation efforts are designed and implemented. While abiotic effects of the grazing deterrents on coral mortality and growth were not measured in this study, they were likely negligible. The nails used as grazing deterrents had a minimal profile and were unlikely to decrease water flow or light, nor did they cause any observable algal accumulation on surrounding benthos as they allowed access to intermediate- and micro-herbivores (Carpenter 1986; Altman-Kurosaki et al. 2018).

Fish grazing on newly outplanted corals has the potential to jeopardise the success of coral restoration efforts. Fish grazing on recent transplants has been widely reported (Miller and Hay 2010; Horoszowski-Fridman et al. 2015; Page et al. 2018; Koval et al. 2020; Smith et al. 2021). Newly transplanted corals might be more palatable to fish grazers due to vulnerability caused by physiological stress. Stress from transplantation and acclimatisation to the new harsher reef environment (e.g., higher light and water flow conditions) can reduce the performance of corals after outplant (Forrester et al. 2012). Corals in *ex-situ* nurseries are acclimatised to low light conditions (Gantt et al. 2023), often having darker pigmentation than wild colonies. This darker pigmentation is lost several weeks to months after outplanting as the transplants acclimatise to their new environment (Fig. 4e&f, Fig. 3d, Horoszowski-Fridman et al. 2015; Page et al. 2018). Another cause of fish grazing on new transplants could be neophilia (i.e., the attraction of grazers to novel objects in the reef) as they explore new food sources. However, continued fish grazing after the first weeks has been observed (Rivas et al. 2021; Knoester et al. 2023; Rule 2023), indicating that fish habituation to novel objects does not reliably preclude predation and incidental damage. Data from our and other studies, including terrestrial reforestation and vegetation restoration reviews, suggest that long-term protection of outplants from grazing and herbivory is beneficial(Redick and Jacobs 2020; Xu et al. 2023).

Some colonies were grazed to small sizes during the first week and did not recover beyond their initial size at outplant (Fig. 3a). Small colonies generally have lower survivorship than larger ones (Raymundo and Maypa 2004; Vermeij and Sandin 2008; Ligson et al. 2022). This bottleneck can extend until corals reach a size threshold, after which mortality stabilises (Babcock and Mundy 1996; Guest et al. 2014). If grazing is spatially random, a lower probability of grazing on smaller colonies is expected, but given the size of the disturbance from parrotfish bites, it is more likely to encompass the entire colony of smaller rather than larger corals (Hughes and Jackson 1985). Hence, small colonies are more likely to experience whole-colony mortality following direct damage (Bak and Engel 1979). In addition, small remnants of grazed colonies may have insufficient energy reserves for wound healing and defence against competing benthic organisms (Connell 1973; Oren et al. 2001).

Successful coral reef recovery or active restoration both require corals to pass the juvenile survivorship bottleneck. Active restoration may be increasingly important in maintaining ecosystem function with continuing coral reef degradation (Knowlton et al. 2021; Suggett et al. 2024). Large grazers such as parrotfish are essential for coral reef resilience, but their detrimental effect on juvenile corals and coral outplants hinders the success of coral reef restoration efforts. Grazing deterrents can help overcome this bottleneck in areas with larger grazing fish populations. On reefs where there are few large grazers (e.g., due to overfishing), there may be fewer benefits from grazing deterrents, or they may not be needed at all. Although manually adding deterrents to outplanted corals using the approach in this study would not be realistic for large-scale restoration efforts due to the dive time required for installation, it could provide a cheap and accessible method to test if grazing deterrents are required during the design stages of coral transplantation efforts. Coral settlement substrates could be designed with integrated grazing deterrents, particularly as our results showed that even deterrents with reduced height are effective. Alternatively, a modular grazing deterrent could be designed to be deployed at outplant, and if grazing pressure is alleviated several weeks or months later, retrieved and reused. Grazing deterrents can, as demonstrated here, increase juvenile survivorship and should be considered as part of coral restoration strategies using sexually propagated corals.

## Acknowledgements

We want to thank the following for making this research possible. The Palau International Coral Reef Center and all their staff for their unwavering support, including Arius Merep, for help in building the spawning setup. Boat operator Nelson Masang and volunteers and research assistants Kailey Gabrian-Voorhees, Jayven Tachibelmel, Ernesto Andres JR, Leah Marie Bukurou, Craig Johnson and Sharon Koen, who helped with outplanting and collecting data in the field. We also thank the Bureau of Marine Resources for donating the rabbitfish used in this study. The work presented here was funded by the European Research Council Horizon 2020 project CORALASSIST (Project number 725848) awarded to JRG, and the Natural Environment Research Council’s ONE Planet Doctoral Training Partnership Studentship (NE/S007512/1) to LL.

