## Supplements for "Excluding Large Grazers Dramatically Improves Survival of Outplanted Juvenile Corals"

**Coral rearing specifics**

Gamete bundles were collected from the surface with plastic cups and placed into a 100 L tank with 0.2 µm UV-treated FSW. The water was gently stirred to break the remaining bundles, after which they were left undisturbed for 45 minutes to allow for optimal fertilisation. After stirring, six 20-ml aliquots were taken and counted; these showed that approximately 874,000 eggs had been released. After fertilisation, an additional six 20-ml aliquots were collected and counted; these showed an achieved fertilisation rate of 94.6 ± 0.8 % (mean ± SE). The fertilised embryos were skimmed off the surface and added to a 100 L tank with 0.2 µm UV-treated FSW to wash off the sperm, a process that was repeated twice. Embryos were then added to two 45 L tanks at a density of one larva per 3 ml; the surplus was returned to the ocean. After spawning, parental colonies were donated to the Palau Aquarium and displayed in the Reef Crest exhibit. Starting 20 hours after fertilisation, 50 % water changes were performed twice daily using 100 µm filters to siphon out water without disturbing the larvae, after which tanks were topped up with 0.2 µm UV-treated FSW. Gentle aeration through rigid air lines was introduced 36 hours after fertilisation when larvae became motile. Four days after fertilisation, mean larval survivorship was 77.0 ± 7.9 %, assessed using eight 50 ml aliquots. During settlement, twice-daily 50 % water changes were performed in each settlement tank, and aeration was added to maintain water movement. After one week, SUs were divided among four shallow 184 L *ex-situ* nursery tanks with ~4 L/min flow-through 50 µm FSW. Each tank had two aquarium lights (Reef Brite, 48” 50/50 white and blue XHO LED; 200 µmol photons $m^{-2}$ $s^{-1}$ over a 12:12 h diurnal cycle) and two pumps (Hydor Koralia Nano 425 Circulation Pump/Powerhead) to ensure water circulation.

**Table 1** Statistical comparison of mean survival times (days) for juvenile coral colonies outplanted to the reef at six months old with five different levels of grazing deterrents monitored for 14 months (425 days).

| Treatment | Mean survival time (days) | 95% Confidence Interval | |
|---|---|---|---|
| | | Lower bound | Upper bound |
| 4 long nails | 374 | 338.1 | 409.9 |
| 4 short nails | 354 | 319.5 | 388.5 |
| 2 long nails | 289 | 240.0 | 338.0 |
| 2 short nails | 324 | 277.7 | 370.3 |
| Control | 212 | 157.5 | 266.5 |

Pairwise comparisons using Log-Rank test
Bonferroni *P* value adjustment

| Treatment | 4 long | 4 short | 2 long | 2 long |
|---|---|---|---|---|
| 4 short nails | 1.000 | - | - | - |
| 2 long nails | **0.042** | 0.760 | - | - |
| 2 long nails | 1.000 | 1.000 | 1.000 | - |
| Control | **<0.001** | **0.005** | 0.744 | **0.035** |

**Table 2** Grazing events recorded in the video assays. Time is the time of camera deployment on the reef, size is the size of the observed fish to the nearest 5 cm, and bites are the number of bites taken by a single fish. The maturation phases are initial phase (IP) and juvenile phase (JP).

| # | Date | Time | Plot | Treatment | Species | Group | Size | Bites | Phase |
|---|---|---|---|---|---|---|---|---|---|
| 1 | 16/4/22 | 10:47 | 2 | 2 long | *Plectroglyphidodon dickii* | Damselfish | 5 | 1 | |
| 2 | 16/4/22 | 10:47 | 2 | 2 long | *Chlorurus sordidus* | Parrotfish | 15 | 3 | IP/JP |
| 3 | 16/4/22 | 10:47 | 2 | 2 long | *Chlorurus sordidus* | Parrotfish | 15 | 2 | IP/JP |
| 4 | 16/4/22 | 10:47 | 2 | 2 long | *Chlorurus sordidus* | Parrotfish | 15 | 5 | IP/JP |
| 5 | 16/4/22 | 10:47 | 2 | 2 long | *Ctenochaetus striatus* | Surgeonfish | 15 | 6 | |
| 6 | 16/4/22 | 10:47 | 2 | 2 short | *Chlorurus sordidus* | Parrotfish | 15 | 2 | IP/JP |
| 7 | 16/4/22 | 10:47 | 2 | 2 short | *Chlorurus sordidus* | Parrotfish | 15 | 5 | IP/JP |
| 8 | 16/4/22 | 10:47 | 2 | 2 short | *Chlorurus sordidus* | Parrotfish | 15 | 1 | IP/JP |
| 9 | 16/4/22 | 10:47 | 2 | 4 long | *Chlorurus sordidus* | Parrotfish | 15 | 1 | IP/JP |
| 10 | 16/4/22 | 10:47 | 2 | 4 long | *Chlorurus sordidus* | Parrotfish | 15 | 2 | IP/JP |
| 11 | 16/4/22 | 10:47 | 2 | 4 short | *Chlorurus sordidus* | Parrotfish | 15 | 5 | IP/JP |
| 12 | 16/4/22 | 10:47 | 2 | Control | *Chlorurus sordidus* | Parrotfish | 20 | 1 | IP/JP |
| 13 | 16/4/22 | 10:47 | 2 | Control | *Chlorurus sordidus* | Parrotfish | 15 | 8 | IP/JP |
| 14 | 16/4/22 | 10:47 | 2 | Control | *Chlorurus sordidus* | Parrotfish | 15 | 6 | IP/JP |
| 15 | 16/4/22 | 10:47 | 2 | Control | *Chlorurus sordidus* | Parrotfish | 15 | 3 | IP/JP |
| 16 | 16/4/22 | 10:47 | 2 | Control | *Chlorurus sordidus* | Parrotfish | 15 | 1 | IP/JP |
| 17 | 16/4/22 | 10:47 | 2 | Control | *Chlorurus sordidus* | Parrotfish | 15 | 2 | IP/JP |
| 18 | 16/4/22 | 10:47 | 2 | Control | *Chlorurus sordidus* | Parrotfish | 15 | 1 | IP/JP |
| 19 | 17/4/22 | 09:44 | 1 | Control | *Labrichthys unilineatus* | Wrasse | 10 | 2 | |
| 20 | 17/4/22 | 09:44 | 1 | Control | *Labrichthys unilineatus* | Wrasse | 10 | 12 | |
| 21 | 17/4/22 | 09:44 | 1 | Control | *Halichoeres melanurus* | Wrasse | 5 | 1 | IP |
| 22 | 17/4/22 | 09:44 | 1 | Control | *Halichoeres melanurus* | Wrasse | 5 | 2 | |
| 23 | 17/4/22 | 09:28 | 2 | 2 long | *Chlorurus sordidus* | Parrotfish | 10 | 2 | |
| 24 | 17/4/22 | 09:28 | 2 | 2 long | *Chlorurus sordidus* | Parrotfish | 10 | 1 | |
| 25 | 17/4/22 | 09:35 | 3 | 4 long | *Halichoeres hortulanus* | Wrasse | 10 | 1 | |
| 26 | 17/4/22 | 12:16 | 1 | 2 long | *Labrichthys unilineatus* | Wrasse | 10 | 5 | IP |
| 27 | 17/4/22 | 12:16 | 1 | 2 long | *Labrichthys unilineatus* | Wrasse | 10 | 8 | IP |
| 28 | 17/4/22 | 12:16 | 1 | 2 long | *Labrichthys unilineatus* | Wrasse | 10 | 9 | IP |
| 29 | 17/4/22 | 12:16 | 1 | 2 long | *Labrichthys unilineatus* | Wrasse | 10 | 4 | IP |
| 30 | 17/4/22 | 12:16 | 1 | 2 long | *Labrichthys unilineatus* | Wrasse | 10 | 13 | IP |
| 31 | 17/4/22 | 12:16 | 1 | 2 long | *Labrichthys unilineatus* | Wrasse | 10 | 17 | IP |
| 32 | 17/4/22 | 12:16 | 1 | 2 long | *Labrichthys unilineatus* | Wrasse | 10 | 7 | IP |
| 33 | 17/4/22 | 12:16 | 1 | 2 long | *Labrichthys unilineatus* | Wrasse | 10 | 5 | IP |
| 34 | 17/4/22 | 12:16 | 1 | 2 long | *Labrichthys unilineatus* | Wrasse | 10 | 3 | IP |
| 35 | 17/4/22 | 12:16 | 1 | 2 long | *Labrichthys unilineatus* | Wrasse | 10 | 7 | IP |
| 36 | 17/4/22 | 12:16 | 1 | 2 short | *Labrichthys unilineatus* | Wrasse | 10 | 13 | IP |
| 37 | 17/4/22 | 12:16 | 1 | 2 short | *Labrichthys unilineatus* | Wrasse | 10 | 14 | IP |
| 38 | 17/4/22 | 12:16 | 1 | Control | *Labrichthys unilineatus* | Wrasse | 10 | 5 | IP |
| 39 | 17/4/22 | 12:16 | 1 | Control | *Labrichthys unilineatus* | Wrasse | 10 | 7 | IP |
| 40 | 17/4/22 | 12:16 | 1 | Control | *Labrichthys unilineatus* | Wrasse | 10 | 6 | IP |
| 41 | 17/4/22 | 12:16 | 1 | Control | *Labrichthys unilineatus* | Wrasse | 10 | 2 | IP |
| 42 | 17/4/22 | 12:16 | 1 | Control | *Labrichthys unilineatus* | Wrasse | 10 | 7 | IP |
| 43 | 17/4/22 | 12:19 | 2 | 2 long | *Apogon exostigma* | Cardinal fish | 5 | 1 | |
| 44 | 17/4/22 | 12:19 | 2 | 2 short | *Chaetodon citrinellus* | Butterflyfish | 10 | 1 | |

| | | | | | | | | | |
|---|---|---|---|---|---|---|---|---|---|
| 45 | 17/4/22 | 15:10 | 1 | Control | *Labrichthys unilineatus* | Wrasse | 10 | 2 | |
| 46 | 17/4/22 | 15:10 | 1 | Control | *Labrichthys unilineatus* | Wrasse | 10 | 3 | IP |
| 47 | 17/4/22 | 15:06 | 3 | 2 short | *Chaetodon citrinellus* | Butterflyfish | 5 | 2 | |
| 48 | 17/4/22 | 15:06 | 3 | Control | *Balistapus undulatus* | Triggerfish | 15 | 1 | |
| 49 | 18/4/22 | 09:13 | 1 | 2 long | *Ctenochaetus striatus* | Surgeonfish | 10 | 1 | |
| 50 | 18/4/22 | 09:13 | 1 | 2 long | *Labrichthys unilineatus* | Wrasse | 10 | 3 | IP |
| 51 | 18/4/22 | 09:13 | 1 | 4 short | *Chaetodon baronessa* | Butterflyfish | 10 | 2 | |
| 52 | 18/4/22 | 09:09 | 2 | 4 long | *Chlorurus sordidus* | Parrotfish | 15 | 3 | JP |
| 53 | 18/4/22 | 09:04 | 3 | 2 long | *Ctenochaetus striatus* | Surgeonfish | 15 | 1 | |
| 54 | 18/4/22 | 09:04 | 3 | 2 short | *Chaetodon citrinellus* | Butterflyfish | 10 | 2 | |
| 55 | 18/4/22 | 09:04 | 3 | Control | *Thalassoma quinquevittatum* | Wrasse | 5 | 2 | |
| 56 | 18/4/22 | 11:40 | 1 | 2 long | *Labrichthys unilineatus* | Wrasse | 10 | 2 | |
| 57 | 18/4/22 | 11:40 | 1 | 2 long | *Labrichthys unilineatus* | Wrasse | 10 | 6 | |
| 58 | 18/4/22 | 11:40 | 1 | 4 long | *Thalassoma hardwicke* | Wrasse | 5 | 1 | |
| 59 | 18/4/22 | 11:40 | 1 | 4 long | *Thalassoma hardwicke* | Wrasse | 5 | 1 | |
| 60 | 18/4/22 | 11:40 | 1 | Control | *Labrichthys unilineatus* | Wrasse | 10 | 1 | |
| 61 | 18/4/22 | 11:42 | 2 | 2 long | *Halichoeres melanurus* | Wrasse | 5 | 1 | |
| 62 | 18/4/22 | 11:42 | 2 | 2 short | *Chaetodon citrinellus* | Butterflyfish | 10 | 1 | |
| 63 | 18/4/22 | 11:42 | 2 | 4 long | *Halichoeres melanurus* | Wrasse | 5 | 1 | |
| 64 | 18/4/22 | 11:42 | 2 | 4 long | *Chlorurus sordidus* | Parrotfish | 10 | 4 | IP/JP |
| 65 | 18/4/22 | 11:42 | 2 | 4 long | *Halichoeres melanurus* | Wrasse | 5 | 1 | |
| 66 | 18/4/22 | 11:42 | 2 | 4 long | *Chlorurus sordidus* | Parrotfish | 15 | 1 | IP/JP |
| 67 | 18/4/22 | 11:42 | 2 | 4 long | *Halichoeres marginatus* | Wrasse | 10 | 1 | |
| 68 | 18/4/22 | 11:44 | 3 | 2 short | *Ctenochaetus striatus* | Surgeonfish | 10 | 3 | |
| 69 | 18/4/22 | 11:44 | 3 | 4 long | *Scarus psittacus* | Parrotfish | 15 | 2 | IP |
| 70 | 18/4/22 | 11:44 | 3 | 4 long | *Scarus psittacus* | Parrotfish | 15 | 4 | IP |
| 71 | 18/4/22 | 11:44 | 3 | Control | *Scarus psittacus* | Parrotfish | 15 | 1 | IP |
| 72 | 18/4/22 | 14:30 | 1 | 2 long | *Chaetodon baronessa* | Butterflyfish | 10 | 3 | |
| 73 | 18/4/22 | 14:30 | 1 | 2 short | *Chaetodon baronessa* | Butterflyfish | 10 | 2 | |
| 74 | 18/4/22 | 14:30 | 1 | 4 long | *Chaetodon baronessa* | Butterflyfish | 10 | 6 | |
| 75 | 18/4/22 | 14:30 | 1 | Control | *Chaetodon baronessa* | Butterflyfish | 10 | 11 | |
| 76 | 18/4/22 | 14:32 | 2 | 2 long | *Chlorurus sordidus* | Parrotfish | 10 | 3 | |
| 77 | 18/4/22 | 14:32 | 2 | Control | *Halichoeres melanurus* | Wrasse | 5 | 2 | |
